# Comparison of *tet*(X4)-containing contigs from metagenomic sequencing data with plasmid sequences of isolates from a cohort of healthy subjects

**DOI:** 10.1101/2023.11.08.566264

**Authors:** Yichen Ding, Shuan Er, Abel Tan, Jean-Sebastien Gounot, Woei-Yuh Saw, Linda Wei Lin Tan, Yik Ying Teo, Niranjan Nagarajan, Henning Seedorf

**Author notes:** contributed equally (Yichen Ding, Shuan Er).

## Abstract

The recently discovered tigecycline-inactivating enzyme Tet(X4) can confer high-level tigecycline resistance on its hosts, which makes it a public health concern. This study focused on detection, analysis, and characterization of Tet(X4)-positive Enterobacteriaceae from the gut microbiota of a healthy cohort of individuals in Singapore using cultivation-dependent and cultivation-independent approaches. Twelve Tet(X4)-positive Enterobacteriaceae strains that were previously obtained from the cohort were fully genome-sequenced and comparatively analysed. A metagenomic sequencing (MS) dataset of the same samples was mined for contigs that harboured the *tet*(X4) resistance gene. The sequences of *tet*(X4)-containing contigs and plasmids sequences were compared. The presence of the resistance genes *floR* and *catD* (also annotated as *estT*) was detected in the same cassette in 10 and 12 out of the 12 *tet*(X4)-carrying plasmids, respectively. MS detected *tet*(X4)-containing contigs in two out of 109 subjects, while cultivation-dependent analysis previously reported a prevalence of 10.1%. Contig sequences are relatively short (∼14-33 kb) but show high similarity to the respective plasmid sequences of the isolates. The frequent co-occurrence of *florR* and *catD* with *tet*(X4) corroborates the hypothesis that the transmission of *tet*(X4) may have originated from the veterinary sector. Our findings show that MS can complement efforts in the surveillance of antibiotic resistance genes for clinical samples, while it has a lower sensitivity than a cultivation-based method when the target organism have a low abundance. Further optimisation is required if MS is to be utilised in antibiotic resistance surveillance.

## Introduction

Genomic surveillance is one important measure to monitor and control the spread of multidrug resistant bacteria (MDR) in the human population. In general, the prevalence for MDR bacteria is determined through a cultivation-dependent approach, i.e. clinical or environmental samples are inoculated into selective agar plates supplemented with antibiotics to obtain MDR bacterial colonies, which will be confirmed with phenotypic and molecular assays. Selected colonies can be further subjected to whole genome sequencing and comparative genomic analysis to determine their antibiotic resistance genes, plasmid sequences and possible transmission clusters. This approach has allowed detailed insights into the genomic structure of many MDR bacteria by numerous studies, especially for the recently emerged *tet*(X) family-mediated tigecycline-resistant Enterobacteriaceae (1, 2). On the other hand, shotgun metagenomic sequencing characterises the microbial communities in clinical and environmental samples via an unbiased culture-independent approach, in which the total DNA of the samples is extracted and sequenced. Subsequently, the antibiotic resistance genes of interest can be further analysed using the contigs assembled from metagenomic sequencing data to determine their presence in the samples, as well as the associated plasmid types and host species. Shotgun metagenomic sequencing may therefore complement culture-based whole genome sequencing approaches for rapid identification of MDR bacteria, especially when cultivation of bacteria is difficult, or a high-throughput screen is required.

The recently identified *tet*(X) family tigecycline resistance genes could confer high-level resistance to last-resort antibiotics such as tigecycline and eravacycline, which poses a serious threat to public health (3). Among the various emerging *tet*(X) variants, *tet*(X4) has been identified in animals, healthy individuals, and patients in multiple provinces of China and other regions (4), and its successful transmission could be attributed to conjugative plasmids and *IS*CR2-mediated transposition (5). We previously reported that the prevalence of Tet(X4)-producing Enterobacteriaceae in the gut microbiota of healthy individuals in Singapore is 10.1%, and analysed the sequences of two IncI1-type plasmids p2EC1-1 and p94EC-2 that carry *tet*(X4) (1). Here, we further sequenced and characterised additional 12 tigecycline resistant Enterobacteriaceae strains isolated from human faecal samples in Singapore. We show that *tet*(X4) is associated with diverse range of plasmid types and hosts and is potentially co-transferred with florfenicol resistance gene *floR* and tylosin resistance gene *catD*. The latter has only recently been characterized as a serine-dependent macrolide esterase (6). We further leveraged on recently published high-quality metagenomic sequence data for the same faecal samples collected from the cohort to evaluate if contigs assembled from metagenomic sequencing data could reveal *tet*(X4) plasmid sequences (1, 7). Our findings suggest that metagenomic sequencing could complement culture-based surveillance for MDR bacteria if they are present at high abundance in clinical samples.

## Methods

### Ethics

This study was conducted in compliance with the Declaration of Helsinki and national and institutional standards. The collection of samples for this study was approved under the National University of Singapore IRB code H-17-026.

### Sample collection and DNA extraction

The collection and DNA extraction of faecal samples have been described previously (1). In brief, faeces from 109 individuals aged 48 to 76 years old of the Singapore Integrative Omics Study were collected in 2018 using a BioCollector (BioCollective) kit, according to the manufacturer’s instructions. Faecal samples were handled in a Coy anaerobic chamber containing N_2_ (75%), CO_2_ (20%), and H_2_ (5%) gas mixture. Homogenized samples were transferred to 50 mL screw-cap tubes prior to storage at –80°C. The QIAamp Power Faecal Pro DNA kit was used to extract gDNA for genomic (2 × 2 mL pure culture; OD_600_ = 0.17) and metagenomic (faecal material; ∼0.5 g) sequencing. DNA for genomic sequencing was further purified using a Qiagen Genomic-tip 20/G kit as described in the manufacturer’s protocol (Qiagen, Germany). Cells from cultures were concentrated at 10,000 × g for 15 min before DNA extraction. DNA was quantified using a Qubit 1.0 fluorometer with a broad range assay kit (Life Technologies) and a NanoDrop-2000 (Thermo Fisher Scientific).

### Genome sequencing and data analysis

Genomic DNA of previously isolated strains was extracted using Qiagen Genomic-tip 20/G as per manufacturer’s instructions. Whole-genome sequencing was performed using MinION and Illumina Novaseq, followed by genome assembly and polishing using Flye v2.9 (8, 9) and Pilon v1.24 (10), respectively. The assembled complete genomes were subjected to sequence typing by online MLST 2.0 (11), phylogenetic analysis using the Harvest Suite (12), antibiotic resistance gene prediction by ResFinder v4.1 (13), plasmid typing by PlasmidFinder v2.0 (14), and identification of insertion sequences by ISFinder (15). Comparative sequence analysis was performed using EasyFig v2.2.5 (16) running BLAST+ 2.13.0 (17).

### Metagenomic sequencing assembly and analysis

MS contigs are derived from the Singapore Platinum Metagenomes Project (SPMP) (7) which was conducted on DNA extracted from the same fecal samples that were also used for the cultivation-based analysis. Contigs containing the *tet*(X4) gene were identified using BLAST and subsequent verification using ResFinder with default settings (13, 17).

### Colony forming unit (CFU) counting

CFU counting experiment was done for our previous study (1). Briefly, frozen faecal samples were weighed and inoculated into LB broth, followed by incubation at 37°C with 200 rpm shaking for 3 h. The faecal suspensions were then serially diluted in 0.9% NaCl and spotted onto MacConkey agar plates supplemented with 2 mg/L eravacycline dihydrochloride. The CFU were enumerated after incubation at 37°C for 18 h, and the results were normalised to CFU per gram of input faecal sample.

### GenBank accession numbers

MS short and long reads can be found under BioProject number PRJEB49168, genomes sequences under BioProject number PRJNA599529.

## Results and discussion

### Characterization of *tet*(X4)-carrying plasmids by whole genome sequencing

Twelve Enterobacteriaceae strains that are positive for *tet*(X4) were previously isolated from human fecal samples on MacConkey agar plates supplemented with eravacycline (1). Their genomes have been sequenced to complete-genome level by Illumina and Nanopore. In total, *tet*(X4) was carried by 7 different plasmid types, including IncHI1A/B-IncFIA (n=3), IncFIB (n=2), IncI (Gamma, n=1), IncX1 (n=1), IncFIA/B-IncI (n=1), IncFII (n=1), IncR (n=1), while two plasmids are non-typable (Figure 1a). The host bacterial species include *E. coli* (n=10), *K. pneumoniae* (n=1, isolate 64EVAM), and *E. cloacae* (n=1, isolate 53EVA) (Figure 1a). In particular, the ten *tet*(X4)-positive *E. coli* strains belonged to ten different sequence types (Figure 1a). These results suggested a broad range of *E. coli* strains with diverse genetic background had been associated with *tet*(X4) in Singapore, which is consistent with findings previously reported in China (5).

**Figure 1.**
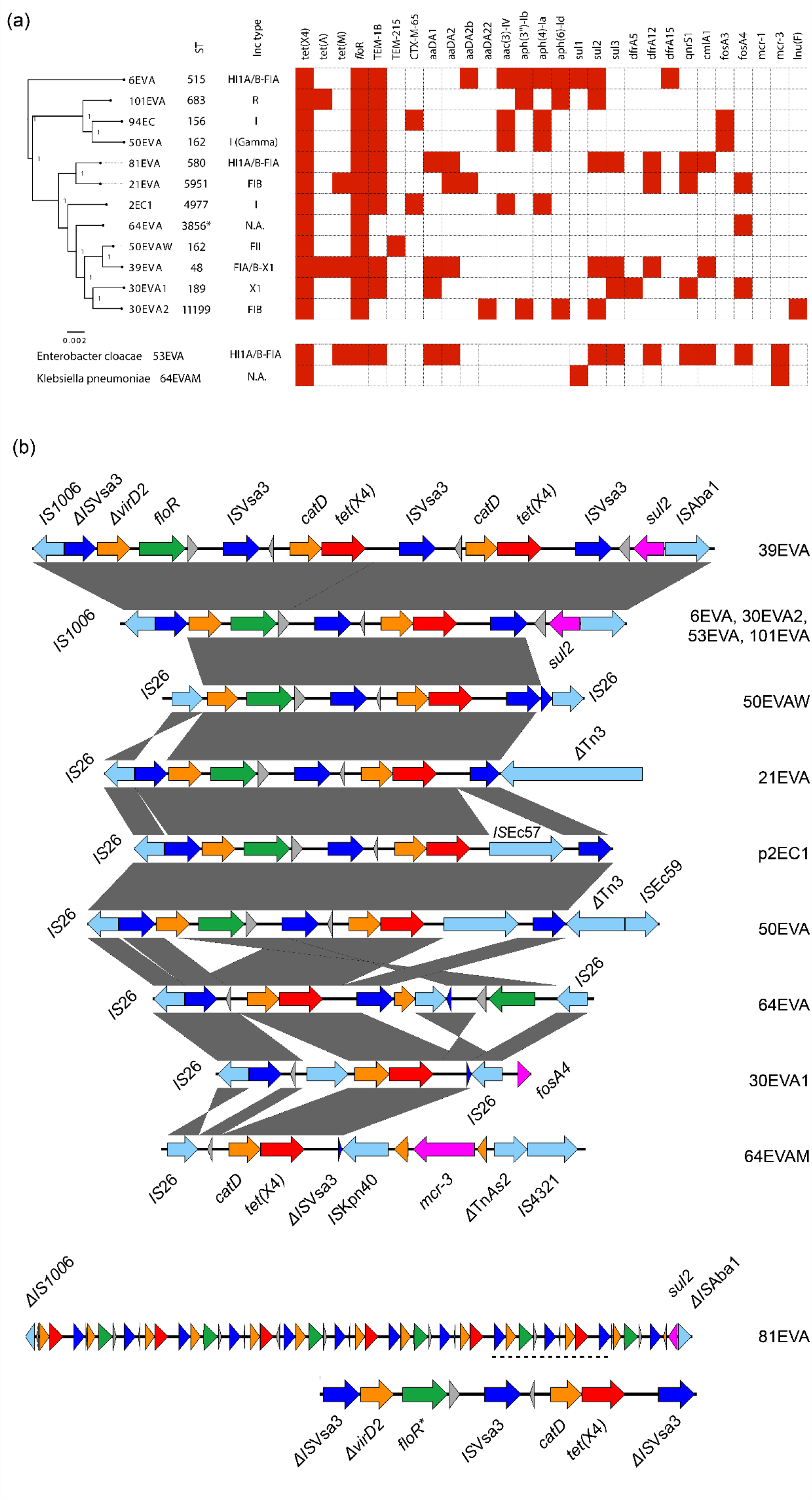
**(a)** Antibiotic resistance gene profiling and phylogenetic analyses of *tet*(X4)-positive Enterobacteriaceae strains. The phylogeny, sequence type and incompatibility group of the *tet*(X4)-harbouring plasmids are shown in the figure (N.A. indicates the plasmid is not typeable). The heat map shows the antibiotic resistance genes carried by *tet*(X4)-carrying plasmid. The presence of antibiotic resistance genes are indicated by red blocks. **(b)** Comparison of *tet*(X4) genetic environments. Open reading frames and their directions of transcription are indicated by colored arrows. Red: *tet*(X4); blue: *IS*Vsa3 (corresponding to *IS*CR2); green: *floR*; magenta: other antibiotic resistance genes; light blue: mobile genetic elements and transposases; orange: genes with putative functions; grey: hypothetical proteins. Truncated genes are indicated by symbol Δ, whereas *flo*R contains a mutation is indicated by an asterisk (*) for isolate 81EVA.

We further characterized and compared the genetic environments of *tet*(X4) among the ten plasmids (Figure 1b). We also included two previously reported isolates 2EC1 and 94EC, which carry *tet*(X4) and *bla*_CTX-M-65_ in our comparative genomic analysis (1). It was found that *tet*(X4) is closely associated with *IS*Vsa3 by having at least one copy of *IS*Vsa3 at its upstream, except for 64EVAM. This is consistent with previous studies showing *tet*(X4) is probably mobilized via *IS*Vsa3 (*IS*CR2)-mediated transposition (1, 2). In addition, we found that other resistance genes are co-occurring with *tet(X4)*, i.e. the *catD* is found in all 12 cassettes, while florfenicol resistance gene *floR* is located in the same cassette in 10 out of the 12 *tet*(X4)-carrying plasmids (Figure 1b), including Δ*IS*Vsa3-Δ*virD2*-*floR*-*IS*Vsa3-*catD*-*tet*(X4)-*IS*Vsa3 (39EVA, 6EVA, 30EVA2, 53EVA, 101EVA, and 81EVA), Δ*IS*Vsa3-Δ*virD2*-*floR*-*IS*Vsa3-*catD*-*tet*(X4)-*IS*EC57-*IS*Vsa3 (50EVA), *IS*26-Δ*virD2*-*floR*-*IS*Vsa3-*catD*-*tet*(X4)-*IS*Vsa3 (21EVA, 50EVAW), and *IS*26-*IS*Vsa3-*catD*-*tet*(X4)-*IS*Vsa3-orf-*IS*26-*floR*-*IS*26 (64EVA). For 30EVA1 and 64EVAM, although *floR* and *tet*(X4) are not in the same cassette, *floR* was either on the same plasmid as *tet*(X4) in 30EVA1, or carried by another plasmid in 64EVAM (Figure 1a). Such close association was not found for other antibiotic resistance genes identified in the 12 strains (Figure 1). The *floR* gene could confer resistance to florfenicol and chloramphenicol, while *catD* confers resistance against 16 membered ring-containing macrolide antibiotics including tylosin, tilmicosin and tildipirosin (18-20). Most of these antibiotics are commonly used as veterinary medicine in aquaculture (21). Similarly, the emergence of *tet*(X4) was suggested to be related to the overuse of tetracycline in food industry (2). The co-carriage of *floR, catD*, and *tet*(X4) by MDR plasmids isolated in healthy individuals in Singapore suggested that their origin might be linked to animal husbandry in the region, while their transmission might be mediated by animal products as suggested by previous studies (22, 23).

### Evaluation of shotgun metagenomic sequencing in detection of *tet*(X4)-carrying plasmids

In total, 11 faecal samples contain *tet*(X4)-positive Enterobacteriaceae, and the *tet*(X4)-carrying plasmid sequences were analysed in this study (Figure 1) and in our previous study (1). To assess if shotgun metagenomic sequencing could detect *tet*(X4)-carrying plasmids, we further screened the contigs assembled from shotgun metagenomic sequencing for *tet*(X4). Interestingly, we found that *tet*(X4)-harbouring contigs can only be detected in two faecal samples (subject SPMP-39 and SPMP-94). The sizes of the *tet*(X4)-harbouring contigs (14-33 kbp) were shorter than the plasmids (101-134 kbp). A comparison of the *tet*(X4)-harbouring contigs with the plasmid sequences revealed high homology of the contigs to the plasmid sequences (Figure 2). This finding indicates that shotgun metagenomic sequencing may potentially aid in the detection of *tet*(X4) and its surrounding genetic environment.

**Figure 2.**
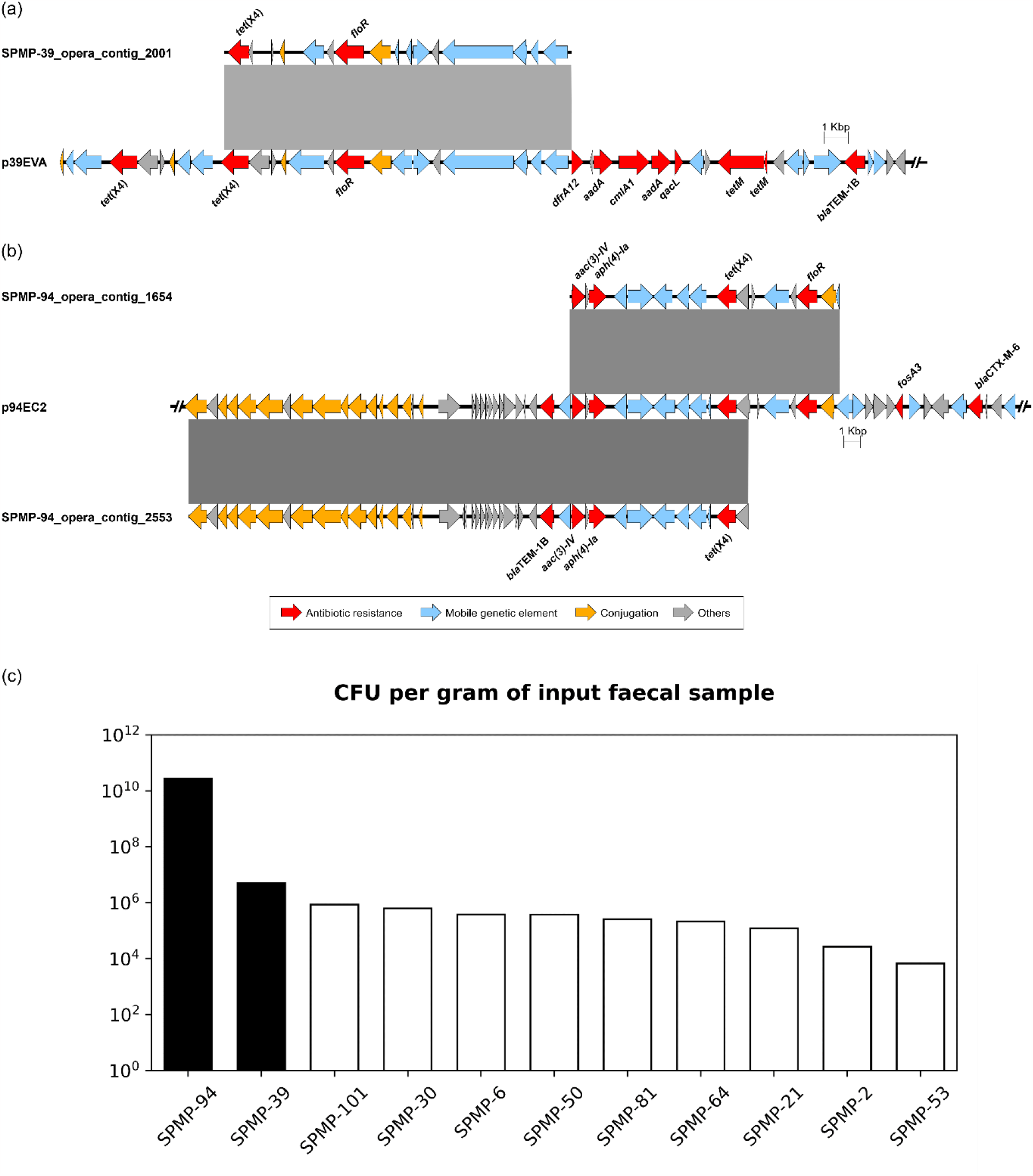
Comparative analysis between metagenomic sequencing-assembled contigs and plasmid sequences containing *tet*(X4) genes derived from **(a)** SPMP-39 and **(b)** SPMP-94. Genes and their respective transcriptional directions are represented by arrows with colours indicating their functional classifications. Shown are truncated sequences of the plasmids. The grey shaded rectangles indicate regions (>4500 bp) of **(a)** 98.8% and **(b)** >99.5% homology. **(c)** Colony forming unit counts of *tet*(X4)-positive Enterobacteriaceae from faecal samples. Contigs assembled from shotgun metagenomic sequencing data contained *tet*(X4)-carrying contigs for subjects SPMP-39 and SPMP-94 (black bars) but not for the other faecal samples (white bars). Of note, the faecal samples were incubated at in LB broth prior to inoculation onto selective agar plate (Methods), this is to allow the *tet*(X4)-positive Enterobacteriaceae to recover from frozen stock before being exposed to the antibiotic for accurate CFU counting. All faecal samples were incubated under the same conditions for the same period of time.

Enterobacteriaceae is often present in low abundance in the human gut, which may potentially result in lower sensitivity for the detection of its associated antibiotic resistance genes when using shotgun metagenomic sequencing. We therefore wondered if the detection of *tet*(X4)-carrying contigs from shotgun metagenomic sequencing data is related to the abundance of the *tet*(X4)-positive Enterobacteriaceae in the faecal samples. Interestingly, out of the three *tet*(X4)-harbouring contigs identified, two of them were detected in subject SPMP-94 who uncoincidentally has a much higher CFU count – by four orders of magnitude – than subject SPMP-39 (Figure 2c). Thus, these results suggest that shotgun metagenomic sequencing could detect *tet*(X4)-harbouring plasmids when the bacteria containing the plasmid is present in high abundance in clinical samples.

Taken together, we report that *tet*(X4) is associated with a broad range of plasmids and host bacteria in the gut of healthy Singaporeans and is closely associated with florfenicol resistance gene *floR and* tylosin resistance gene *catD*. By comparing the contigs assembled from shotgun metagenomic sequencing, we show that this approach could complement culture-based detection of *tet*(X4) plasmids in human faecal samples when present at higher abundance. Further optimisation is required if metagenomic sequencing should be used to discover MDR from clinical and environmental samples. However, selective cultivation currently remains the most reliable and cost-effective approach for detection of antibiotic resistant bacteria.

## Transparency declarations

None to declare.

